# mRNA vaccine-elicited antibodies to SARS-CoV-2 and circulating variants

**DOI:** 10.1101/2021.01.15.426911

**Authors:** Zijun Wang, Fabian Schmidt, Yiska Weisblum, Frauke Muecksch, Christopher O. Barnes, Shlomo Finkin, Dennis Schaefer-Babajew, Melissa Cipolla, Christian Gaebler, Jenna A. Lieberman, Thiago Y. Oliveira, Zhi Yang, Morgan E. Abernathy, Kathryn E. Huey-Tubman, Arlene Hurley, Martina Turroja, Kamille A. West, Kristie Gordon, Katrina G. Millard, Victor Ramos, Justin Da Silva, Jianliang Xu, Robert A. Colbert, Roshni Patel, Juan Dizon, Cecille Unson-O’Brien, Irina Shimeliovich, Anna Gazumyan, Marina Caskey, Pamela J. Bjorkman, Rafael Casellas, Theodora Hatziioannou, Paul D. Bieniasz, Michel C. Nussenzweig

## Abstract

To date severe acute respiratory syndrome coronavirus-2 (SARS-CoV-2) has infected over 100 million individuals resulting in over two million deaths. Many vaccines are being deployed to prevent coronavirus disease 2019 (COVID-19) including two novel mRNA-based vaccines^1,2^. These vaccines elicit neutralizing antibodies and appear to be safe and effective, but the precise nature of the elicited antibodies is not known^3–6^. Here we report on the antibody and memory B cell responses in a cohort of 20 volunteers who received either the Moderna (mRNA-1273) or Pfizer-BioNTech (BNT162b2) vaccines. Consistent with prior reports, 8 weeks after the second vaccine injection volunteers showed high levels of IgM, and IgG anti-SARS-CoV-2 spike protein (S) and receptor binding domain (RBD) binding titers^3,5,6^. Moreover, the plasma neutralizing activity, and the relative numbers of RBD-specific memory B cells were equivalent to individuals who recovered from natural infection^7,8^. However, activity against SARS-CoV-2 variants encoding E484K or N501Y or the K417N:E484K:N501Y combination was reduced by a small but significant margin. Consistent with these findings, vaccine-elicited monoclonal antibodies (mAbs) potently neutralize SARS-CoV-2, targeting a number of different RBD epitopes in common with mAbs isolated from infected donors. Structural analyses of mAbs complexed with S trimer suggest that vaccine- and virus-encoded S adopts similar conformations to induce equivalent anti-RBD antibodies. However, neutralization by 14 of the 17 most potent mAbs tested was reduced or abolished by either K417N, or E484K, or N501Y mutations. Notably, the same mutations were selected when recombinant vesicular stomatitis virus (rVSV)/SARS-CoV-2 S was cultured in the presence of the vaccine elicited mAbs. Taken together the results suggest that the monoclonal antibodies in clinical use should be tested against newly arising variants, and that mRNA vaccines may need to be updated periodically to avoid potential loss of clinical efficacy.

---

Between 19 October 2020 and 15 January 2021, 20 volunteers who received two doses of the Moderna (n=14) or Pfizer-BioNTech mRNA (n=6) vaccines were recruited for blood donation and analyzed. Ages of the analyzed volunteers ranged from 29-69 years (median 43); 12 (60%) were male and 8 (40%) female. 16 participants identified as Caucasian, 2 as Hispanic, and 1 as African American or Asian, respectively. The time from the second vaccination to sample collection varied between 3-14 weeks with an average of 8 weeks. None of the volunteers had a history of prior SARS-CoV-2 infection and none experienced serious adverse events after vaccination (Extended Data Table 1).

## Vaccine plasma binding and neutralizing activity against SARS-CoV-2

Plasma IgM, IgG and IgA responses to SARS-CoV-2 S and RBD were measured by enzyme-linked immunosorbent assay (ELISA)^7,8^. All individuals tested showed reactivity to S and RBD that was significantly higher compared to pre-COVID-19 historic controls (Extended Data Fig 1a-f). As might be expected anti-S and -RBD IgG levels were higher than IgM or IgA. Moreover, there was a strong positive correlation between anti-RBD and anti-S response in all three immunoglobulin isotypes measured (Extended Data Fig 1g-i). In line with previous reports^3,6,9^, IgG and IgM levels were significantly higher in the vaccinated group compared to a cohort of convalescent patients assayed 1.3 and 6.2 months after infection, while IgA levels were similar (Extended Data Fig 1j-l).

Plasma neutralizing activity was determined using human immunodeficiency virus-1 (HIV-1) pseudotyped with SARS-CoV-2 S protein^7,8,10^. In agreement with previous reports^3,6,9^, there was a broad range of plasma neutralizing activity 3-14 weeks after the second vaccine dose that was similar to that elicited by natural infection in a convalescent cohort after 1.3 months, and greater than the activity at 6.2 months after infection (Fig. 1a, Extended Data Table 1). There was no significant difference in neutralizing activity between the Moderna and Pfizer-BioNTech vaccines (Fig. 1b). Whereas convalescent antibody titers tend to correlate with severity and length of time of infection, additional sampling would be required to understand the correlates of the magnitude of the vaccine responses. As expected, plasma neutralizing activity was directly correlated to anti-S and -RBD binding titers in ELISAs^7,8^ (Fig. 1c, d, and Extended Data Fig. 2a-d). Finally, RBD and S binding, and neutralizing activities were directly correlated to the time between the first vaccine dose and blood donation with significantly reduced levels in all 3 measurements with time (Fig. 1e, f and g, and Extended Data Fig. 2e-h)^11^. However, this and other small studies^3,11^ cannot accurately predict the half-life of the neutralizing response. Larger numbers of individuals in diverse cohorts will need to be studied to determine the precise half-life of the vaccine elicited neutralizing response.

**Fig. 1.**
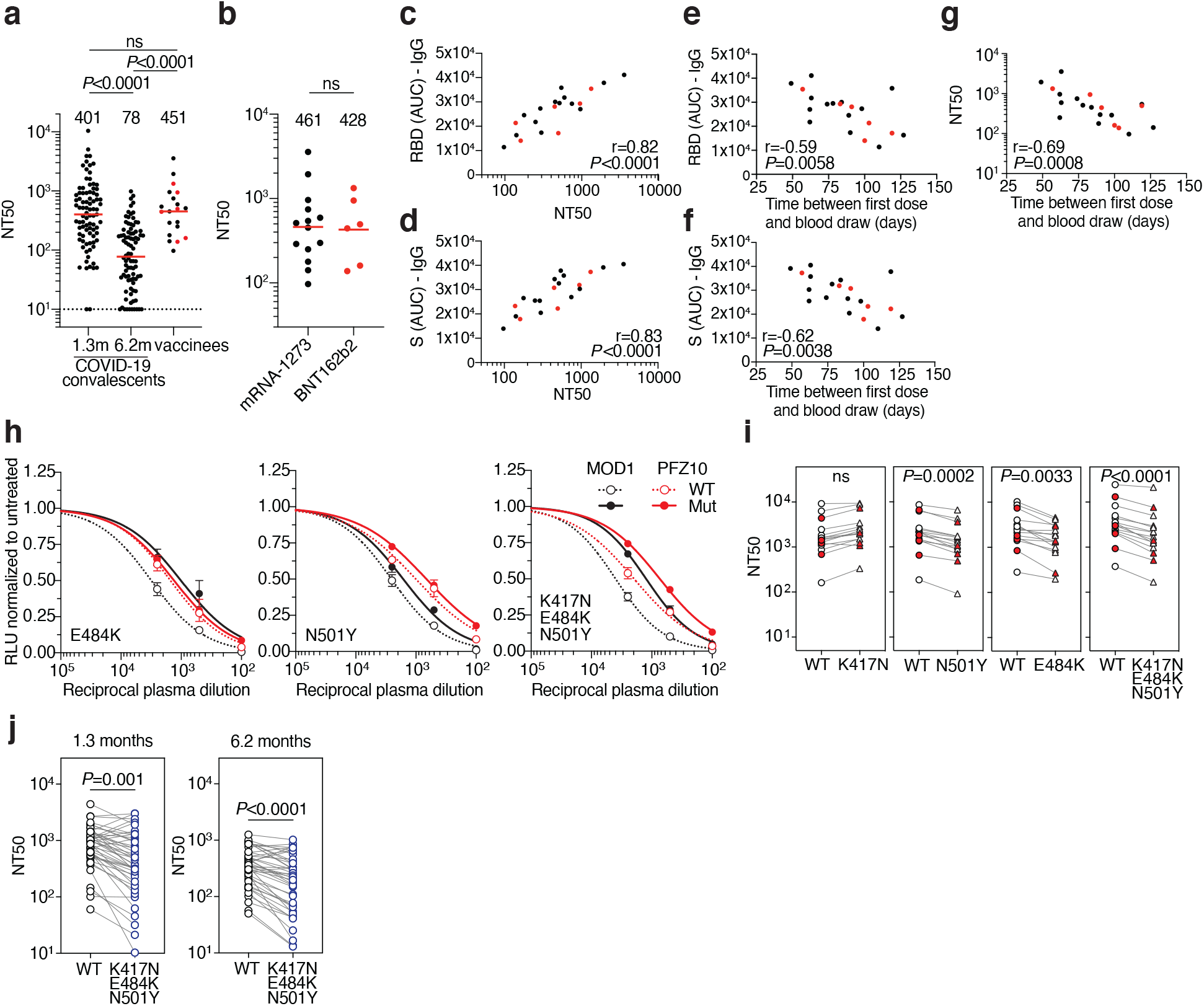
Plasma neutralizing activity. **a,** SARS-CoV-2 pseudovirus neutralization assay. NT_50_ values for COVID-19 convalescent plasma measured at 1.3 months^8^ and 6.2 months^7^ after infection as well as plasma from vaccinees. NT_50_ values lower than 10 were plotted at 10. Mean of 2 independent experiments. Red bars and indicated values represent geometric mean NT_50_ values. Statistical significance was determined using the two-tailed Mann-Whitney U-test. Pre-COVID-19 historical control plasma was analyzed as a negative control and showed no detectable neutralization (NT_50_<10). **b**, NT_50_ values for Moderna mRNA-1273 (black) and Pfizer-BioNTech BNT162b2 (red) vaccine recipients. Red bars and indicated values represent geometric mean NT_50_ values. Statistical significance was determined using the two-tailed Mann-Whitney U-test. **c**, Anti-RBD IgG AUC (Y axis) plotted against NT_50_ (X axis) r=0.82, p<0.0001. **d,** Anti-S IgG AUC (Y axis) plotted against NT_50_ (X axis) r=0.83, p<0.0001. **e**, Anti-RBD IgG AUC (Y axis) plotted against time between first dose and blood draw (X axis) r=−0.59 p=0.0058. **f**, Anti-S IgG AUC (Y axis) plotted against time between first dose and blood draw (X axis) r=−0.62 p=0.0038. **g**, NT_50_ (Y axis) plotted against time between first dose and blood draw (X axis) r=−0.69 p=0.0008. The r and p values for correlations in **c-g** were determined by two-tailed Spearman correlation. Moderna vaccinees are in black and Pfizer-BioNTech in red. **h.** Examples of neutralization assays, comparing the sensitivity of pseudotyped viruses with WT and RBD mutant SARS-CoV-2 S proteins to vaccinee plasma. MOD1 and PFZ10 indicate two representative individuals receiving the Moderna and Pfizer-BioNTech vaccine, respectively (for details see Ext. Data Table 1). **i,** NT50 values for vaccinee plasma (n=15) neutralization of pseudotyped viruses with WT and the indicated RBD-mutant SARS-CoV-2 S proteins. Pfizer-BioNTech vaccinees in red. **j**, NT50 values for convalescent plasma (n=45) neutralization of pseudotyped viruses with WT and KEN (K417N/E484K/N501Y) SARS-CoV-2 S proteins. Statistical significance in **i** and **j** was determined using one tailed t-test. All experiments were performed a minimum of 2 times. Pseutotyped viruses containing the E484K mutation and corresponding WT controls contain the R683G mutation (for details see methods).

To determine whether plasma from vaccinated individuals can neutralize circulating SARS-CoV-2 variants of concern and mutants that arise *in vitro* under antibody pressure ^12,13^, we tested vaccinee plasma against a panel of 10 mutant pseudotype viruses including recently reported N501Y (B1.1.7 variant), K417N, E484K and the combination of these 3 RBD mutations (501Y.V2 variant)^14–19^. Vaccinee plasma was significantly less effective in neutralizing the HIV-1 virus pseudotyped with certain SARS-CoV-2 mutant S proteins (Fig. 1h and i and Extended Data Fig. 2j). Among the volunteer plasmas tested there was a 1- to 3-fold decrease in neutralizing activity against E484K, N501Y and the K417N:E484K:N501Y combination (p= 0.0033, p=0.0002, and p<0.0001, respectively, Fig. 1h and i). Similarly, convalescent plasma obtained 1.3 and 6.2 months after infection was 0.5- to 29- and 0.5- to 20.2-fold less effective in neutralizing the K417N:E484K:N501Y combination (p=0.001 and p<0.0001, respectively, Fig. 1j, Extended Data Table 2). We conclude that the plasma neutralizing activity elicited by either mRNA vaccination or natural infection is variably but significantly less effective against pseudoviruses that carry RBD mutations found in emerging SARS-CoV-2 variants.

## Vaccine-elicited SARS-CoV-2 RBD-specific monoclonal antibodies

Although circulating antibodies derived from plasma cells wane over time, long-lived immune memory can persist in expanded clones of memory B cells^7,20^. We used flow cytometry to enumerate the circulating SARS-CoV-2 RBD-specific memory B cells elicited by mRNA immunization^7,8^ (Fig. 2a, Extended Data Fig. 3a and b). We focused on the RBD since it is the target of the majority of the more potent SARS-CoV-2 neutralizing antibodies discovered to date^21–26^. Notably, the percentage of RBD-binding memory B cells in vaccinees was significantly greater than in naturally infected individuals assayed after 1.3 months, but similar to the same individuals assayed after 6.2 months (Fig. 2b). The percentage of RBD-binding memory B cells in vaccinees was not correlated to the time after vaccination (Extended Data Fig. 3c). Thus, mRNA vaccination elicits a robust SARS-CoV-2 RBD-specific B cell memory response that resembles natural infection.

**Fig. 2.**
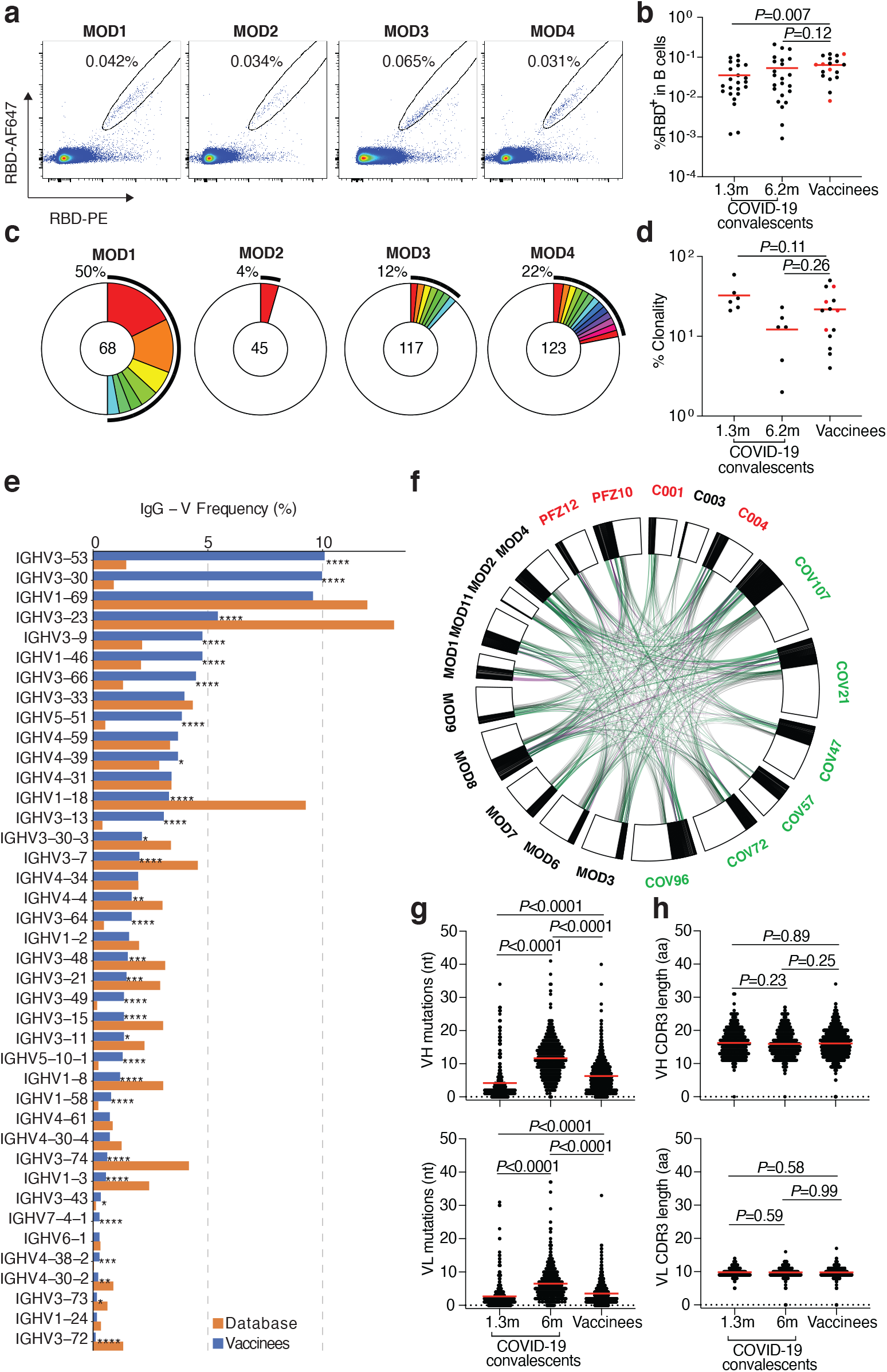
Memory B cell antibodies. **a,** Representative flow cytometry plots showing dual AlexaFluor-647-RBD and PE-RBD binding B cells for 4 vaccinees. **b,** as in **a,** dot plot summarizes the percentage of RBD binding B cells in 19 vaccinees, in comparison to a cohort of infected individuals assayed 1.3 and 6.2 months after infection^7,8^. Individuals who received the Moderna vaccine are shown in black and Pfizer-BioNTech vaccine recipients in red. Red horizontal bars indicate mean values. Statistical significance was determined using two-tailed Mann–Whitney U-tests. **c,** Pie charts show the distribution of antibody sequences from the 4 individuals in **a.** The number in the inner circle indicates the number of sequences analyzed. Pie slice size is proportional to the number of clonally related sequences. The black outline indicates the frequency of clonally expanded sequences. **d,** as in **c,** graph shows relative clonality among 14 vaccinees assayed, individuals who received the Moderna vaccine are shown in black and Pfizer-BioNTech vaccine recipients in red. Red horizontal bars indicate mean values. Statistical significance was determined using two-tailed Mann–Whitney U-tests. **e,** Graph shows relative abundance of human IGVH genes Sequence Read Archive accession SRP010970 (orange), and vaccinees (blue). A two-sided binomial test was used to compare the frequency distributions, significant differences are denoted with stars (* p < 0.05, ** p < 0.01, *** p < 0.001, **** = p < 0.0001). **f,** Clonal relationships between sequences from 14 vaccinated individuals (Moderna in black, Pfizer-BioNTech in red Extended Data Table 3) and naturally infected individuals (in green, from^7,8^). Interconnecting lines indicate the relationship between antibodies that share V and J gene segment sequences at both IGH and IGL. Purple, green and grey lines connect related clones, clones and singles, and singles to each other, respectively. **g,** Number of somatic nucleotide mutations in the IGVH (top) and IGVL (bottom) in vaccinee antibodies (Extended Data Table 3) compared to natural infection obtained 1.3 or 6.2 months after infection^7,8^. Statistical significance was determined using the two-tailed Mann–Whitney U-tests and red horizontal bars indicate mean values. **h,** as in **g,** but for CDR3 length.

To examine the nature of the antibodies produced by memory B cells in response to vaccination, we obtained 1,409 paired antibody heavy and light chains from RBD binding single B cells from 14 individuals (n=10 Moderna and n=4 Pfizer-BioNTech vaccinees) (Extended Data Table 3). Expanded clones of cells comprised 4-50% of the overall RBD binding memory B cell compartment (Fig. 2c and d, and Extended Data Fig. 3d). Similar to natural infection, IGVH 3-53, and 3-30 and some IGVL genes were significantly over-represented in the RBD-binding memory B cell compartment of vaccinated individuals (Fig. 2e, Extended Data Fig. 4a). In addition, antibodies that share the same combination of IGHV and IGLV genes in vaccinees comprised 39% of all the clonal sequences (Extended Data Fig. 4b) and 59% when combined with naturally infected individuals^7,8^ (Fig. 2f), and some of these antibodies were nearly identical (Extended Data Table 3 and 4). The number of V gene nucleotide mutations in vaccinees is greater than in naturally infected individuals assayed after 1.3 months, but lower than that in the same individuals assayed after 6.2 months (Fig. 2g and Extended Data Fig. 5a). The length of the IgH CDR3 was similar in both natural infected individuals and vaccinees and hydrophobicity was below average^27^ (Fig. 2h and Extended Data Fig. 5a and b). Thus, the IgG memory response is similar in individuals receiving the Pfizer-BioNTech and Moderna vaccines and both are rich in recurrent and clonally expanded antibody sequences.

One hundred and twenty-seven representative antibodies from 8 individuals were expressed and tested for reactivity to the RBD (Extended Data Table 5). The antibodies included: (1) 76 that were randomly selected from those that appeared only once, and (2) 51 representatives of expanded clones. Of the antibodies tested 98% (124 out of the 127) bound to RBD indicating that single cell sorting by flow cytometry efficiently identified B cells producing anti-RBD antibodies (Extended Data Fig. 6a-c and Table 5). In anti-RBD ELISAs the mean half-maximal effective concentration (EC_50_) was higher than that observed in infected individuals after 6.2 months but not significantly different from antibodies obtained 1.3 months after infection (Extended Data Fig. 6a, and Table 5 and ^7,8^). To examine memory B cell antibodies for binding to circulating SARS-CoV-2 variants and antibody resistant mutants we performed ELISAs using mutant RBDs^12,15,28–31^. Twenty-two (26%) of the 84 antibodies tested showed at least 5-fold decreased binding to at least one of the mutant RBDs (Extended data Fig. 6d-n and Table 5).

SARS-CoV-2 S pseudotyped viruses were used to measure the neutralizing activity of all 127 antibodies^7,8,10^ (Fig. 3a, Extended Data Table 5). Consistent with the plasma neutralization results, the geometric mean neutralization half-maximal inhibitory concentration of the Pfizer-BioNTech and Moderna vaccinee antibodies were not significantly different from each other or to antibody collections obtained from naturally infected individuals 1.3 or 6.2 months after infection (Fig. 3a and ^7,8^).

**Fig. 3:**
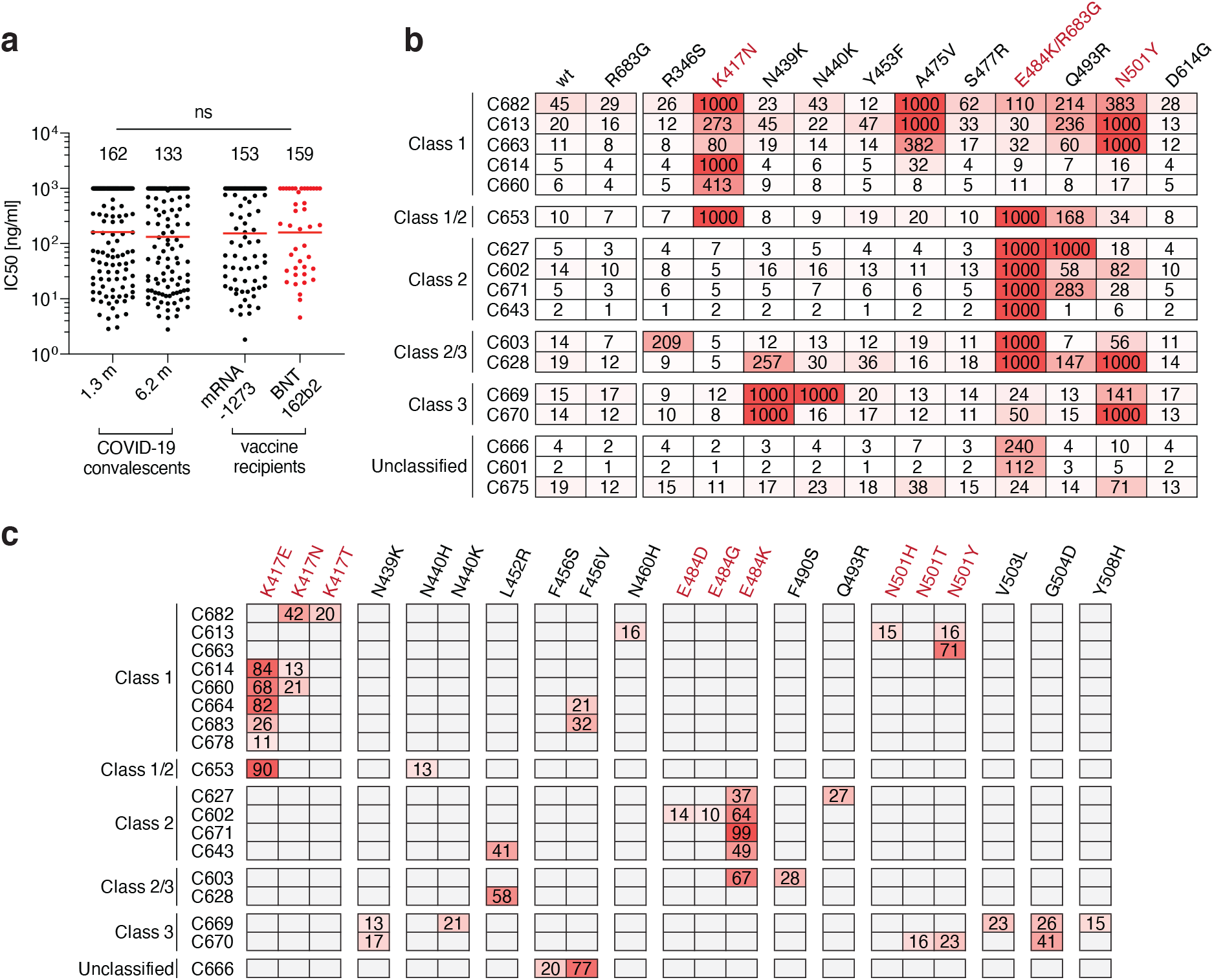
Anti-SARS-CoV-2 RBD monoclonal antibody neutralizing activity. **a,** SARS-CoV-2 pseudovirus neutralization assay. IC_50_ values for antibodies cloned from COVID-19 convalescent patients measured at 1.3 and 6.2 months^7,8^ after infection as well as antibodies cloned from Moderna mRNA-1273 (black) and Pfizer-BioNTech BNT162b2 (red) mRNA-vaccine recipients. Antibodies with IC_50_ values above 1000 ng/ml were plotted at 1000 ng/ml. Mean of 2 independent experiments. Red bars and indicated values represent geometric mean IC_50_ values in ng/ml. Statistical significance was determined using the two-tailed Mann-Whitney U-test. Isotype control antibody was analyzed in parallel and showed no detectable neutralization. **b,** IC_50_ values for 17 selected mAbs for neutralization of wild type and the indicated mutant SARS-CoV-2 pseudoviruses. Color gradient indicates IC_50_ values ranging from 0 (white) to 1000 ng/ml (red). **c**, Antibody selection pressure can drive emergence of S variants in cell culture; the percentage of sequence reads encoding the indicated RBD mutations after a single passage of rVSV/SARS-CoV-2 in the presence of the indicated antibodies is tabulated. Color gradient indicates percentage of sequence reads bearing the indicated mutation ranging from 0 (white) to 100 (red). Positions for which no sequence read was detected are shown in grey. The percentages calculated for a given position are based on all the reads, and not just the reads that include that position. K417N, E484K/R683G and N501 are highlighted in **b** and **c** as they constitute important circulating variants.

To examine the neutralizing breadth of the monoclonal antibodies and begin to map their target epitopes we tested 17 of the most potent antibodies (Extended data Table 6), 8 of which carried IgHV3-53, against a panel of 12 SARS-CoV-2 variants: A475V is resistant to class 1 antibodies (structurally defined as described^29^); E484K and Q493R are resistant to class 2 antibodies^7,8,12,13,29,30,32,33^; while R346S, N439K, and N440K are resistant to class 3 antibodies^7,8,12,13,29,33^. Additionally, K417N, Y453F, S477R, N501Y, and D614G represent circulating variants some of which have been associated with rapidly increasing case numbers^14,15,19,33–35^. Based on their neutralizing activity against the variants, all but 3 of the antibodies were provisionally assigned to a defined antibody class or a combination (Fig. 3b). As seen in natural infection, a majority of the antibodies tested (9/17) were at least ten-fold less effective against pseudotyped viruses carrying the E484K mutation^7,12,29^. In addition, 5 of the antibodies were less potent against K417N and 4 against N501Y by ten-fold or more (Fig. 3b). Similar results were obtained with antibodies being developed for clinical use (REGN10987, REGN10933, COV2-2196, COV2-2130, C135 and C144 (Extended data Fig. 7)). However, antibody combinations remained effective against all of the variants tested confirming the importance of using antibody combinations in the clinic (Extended data Fig. 7). Whether less potent antibodies show similar degrees of sensitivity to these mutations remains to be determined.

To determine whether antibody-imposed selection pressure could also drive the emergence of resistance mutations *in vitro,* we cultured an rVSV/SARS-CoV-2 recombinant virus in the presence of each of 18 neutralizing monoclonal antibodies. All of the tested antibodies selected for RBD mutations. Moreover, in all cases the selected mutations corresponded to residues in the binding sites of their presumptive antibody class (Fig. 3b and c). For example, antibody C627, which was assigned to class 2 based on sensitivity to the E484K mutation, selected for the emergence of the E484K mutation *in vitro* (Fig 3c). Notably, 6 of the antibodies selected for K417N, E or T, 5 selected for E484K and 3 selected for N501Y, T or H, which coincide with mutations present in the circulating B.1.1.17/501Y.V1, B.1.351/501Y.V2 and B1.1.28/501.V3 (P.1) variants that have been associated with rapidly increasing case numbers in particular locales^14,17,18,36^.

## Cryo-EM Mapping of Antibody Epitopes

To further characterize antibody epitopes and mechanisms of neutralization, we characterized seven complexes between mAb Fab fragments and the prefusion, stabilized ectodomain trimer of SARS-CoV-2 S glycoprotein^37^ using single-particle cryo-EM (Fig 4 and Extended Data Table 7). Overall resolutions ranged from 5-8 Å (Extended Data Fig. 8) and coordinates from S trimer and representative Fab crystal structures were fit by rigid body docking into the cryo-EM density maps to provide a general assessment of antibody footprints/RBD epitopes. Fab-S complexes exhibited multiple RBD-binding orientations recognizing either ‘up’/’down’ (Fig 4a-j) or solely ‘up’ (Fig 4k-n) RBD conformations, consistent with structurally defined antibody classes from natural infection (Fig 4o)^29^. The majority of mAbs characterized (6 of 7) recognized epitopes that included RBD residues involved in ACE2 recognition, suggesting a neutralization mechanism that directly blocks ACE2-RBD interactions. Additionally, structurally defined antibody epitopes were consistent with RBD positions that were selected in rVSV/SARS-CoV-2 recombinant virus outgrowth experiments, including residues K417, N439/N440, E484, and N501 (Fig 3c and Fig 4f-j,m,n). Taken together, these data suggest that functionally similar antibodies are raised during vaccination and natural infection, and that the RBDs of spike trimers translated from the mRNA delivered by vaccination adopt both ‘up’ and ‘down’ conformations as observed on structures of trimer ectodomains^29^ and trimers on the surface of SARS-CoV-2 virions^38^.

**Fig. 4.**
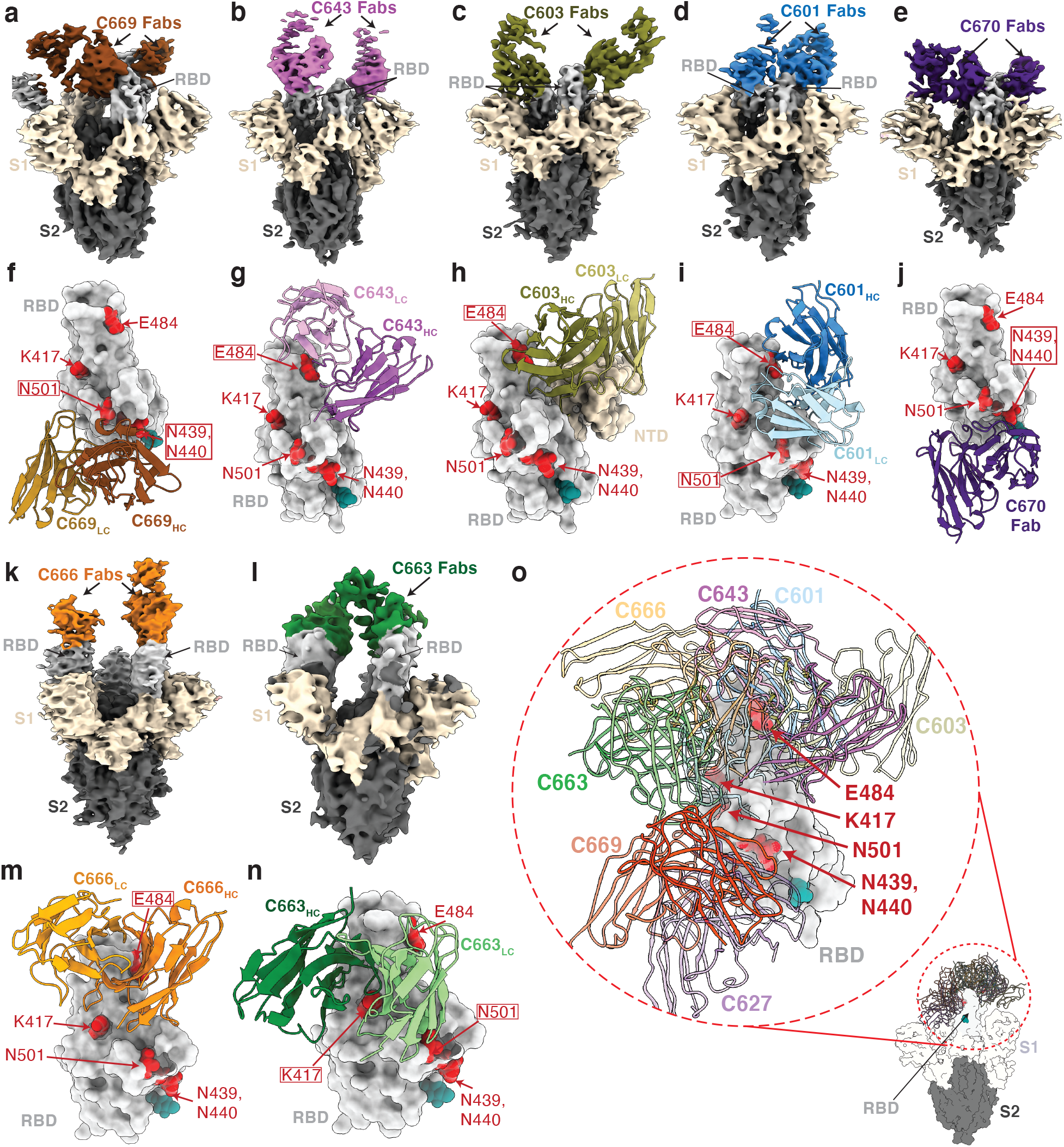
Cryo-EM reconstructions of Fab-S complexes. Cryo-EM densities for Fab-S complexes (**a-e**; **k-l**) and close-up views of antibody footprints on RBDs (**f-j**; **m-n**) are shown for neutralizing mAbs. As expected, due to Fab inter-domain flexibility, cryo-EM densities (**a-e**; **k-l**) were weak for the Fab C_H_-C_L_ domains. Models of antibody footprints on RBDs (**f-j**; **m-n**) are presented as Fab V_H_-V_L_ domains (cartoon) complexed with the RBD (surface). To generate models, coordinates of stabilized S trimer (PDB 6XKL) and representative Fab fragments (PDB 6XCA or 7K8P) with CDR3 loops removed were fit by rigid body docking into the cryo-EM density maps. **a,f,** C669; **b,g,** C643; **c,h,** C603; **d,i,** C601; **e,j,** C670; **k,m,** C666; and **l,n,** C663. RBD residues K417, N439, N440, E484, and N501 are highlighted as red surfaces. The N343 glycan is shown as a teal sphere. **o,** Composite model illustrating targeted epitopes of RBD-specific neutralizing mAbs (shown as V_H_-V_L_ domains in colors from panels **a-l**) elicited from mRNA vaccines.

## Discussion

The mRNA-based SARS-CoV-2 vaccines are safe and effective and being deployed globally to prevent infection and disease. The vaccines elicit antibody responses against the RBD, the major target of neutralizing antibodies^21–26^, in a manner that resembles natural infection. Notably, the neutralizing antibodies produced by mRNA vaccination target the same epitopes as natural infection. The data are consistent with SARS-CoV-2 spike trimers translated from the injected RNA adopting a range of different conformations. Moreover, different individuals immunized with either the Moderna (mRNA-1273) or Pfizer-BioNTech (BNT162b2) vaccines produce closely related and nearly identical antibodies. Whether or not neutralizing antibodies to epitopes other that RBD are elicited by vaccination remains to be determined.

Human neutralizing monoclonal antibodies to the SARS-CoV-2 RBD can be categorized as belonging to 4 different classes based on their target regions on the RBD^29^. Class 1 and 2 antibodies are among the most potent and also the most abundant antibodies^7,8,21,22,25,39^. These antibodies target epitopes that overlap or are closely associated with RBD residues K417, E484 and N501. They are frequently sensitive to mutation in these residues and select for K417N, E484K and N501Y mutations in SARS-CoV-2 S protein expression libraries in yeast and VSV ^12,15,33^. To avert selection and escape, antibody therapies should be composed of combinations of antibodies that target non-overlapping or highly conserved epitopes^8,12,13,33,40–44^.

A number of circulating SARS-CoV-2 variants that have been associated with rapidly increasing case numbers and have particular prevalence in the UK (B.1.1.7/501Y.V1), South Africa (B.1.351/501Y.V2) and Brazil (P.1)^14,17,18,36^. Our experiments indicate that the RBD mutations found in these variants, and potentially others that carry K417N/T, E484K and N501Y mutations, can reduce the neutralization potency of vaccinee and convalescent plasma against SARS-CoV-2 pseudotyped viruses. Although our assays are limited to pseudotyped viruses there is an excellent correlation between pseudotyped and authentic SARS-CoV-2 neutralization assays^10^. In addition, similar results have been reported by others using vaccinee and convalescent plasmas and a variety of different pseudotype and authentic virus assays^15,31,45–49^.

The comparatively modest effects of the mutations on viral sensitivity to plasma reflects the polyclonal nature of the neutralizing antibodies in vaccinee plasma. Nevertheless, emergence of these particular variants is consistent with the dominance of the class 1 and 2 antibody response in infected or vaccinated individuals. We speculate that these mutations emerged in response to immune selection in individuals with non-sterilizing immunity. What the long-term effect of accumulation of mutations on the SARS-CoV-2 pandemic will be is not known, but the common cold coronavirus HCoV-229E evolves antigenic variants that are comparatively resistant to the older sera but remain sensitive to contemporaneous sera^50^. Thus, it is possible that these mutations and others that emerge in individuals with suboptimal or waning immunity will erode the effectiveness of natural and vaccine elicited immunity. The data suggests that SARS-CoV-2 vaccines and antibody therapies may need to be updated and immunity monitored in order to compensate for viral evolution.

## Data reporting

No statistical methods were used to predetermine sample size. The experiments were not randomized and the investigators were not blinded to allocation during experiments and outcome assessment.

## Study participants

To isolate and characterize anti-SARS-CoV-2 RBD antibodies from vaccinees, a cohort of 20 individuals that participated in either the Moderna or Pfizer-BioNTech phase 3 vaccine clinical trials and did not report prior history of SARS-CoV-2 infection was recruited at the NIH Blood Center and the Rockefeller University Hospital for blood donation. Eligible participants included adults, at least 18 years of age with no known heart, lung, kidney disease or bleeding disorders, no history of HIV-1 or malaria infection. All participants were asymptomatic at the time of the study visit and had received a complete 2 dose regimen of either mRNA vaccine. Informed consent was obtained from all participants and the study was conducted in accordance with Good Clinical Practice. The study visits and blood draws were reviewed and approved under the National Institutes of Health’s Federalwide Assurance (FWA00005897), in accordance with Federal regulations 45 CFR 46 and 21 CFR 5 by the NIH Intramural Research Program IRB committee (IRB# 99CC0168, Collection and Distribution of Blood Components from Healthy Donors for In Vitro Research Use) and by the Institutional Review Board of the Rockefeller University (IRB# DRO-1006, Peripheral Blood of Coronavirus Survivors to Identify Virus-Neutralizing Antibodies). For detailed participant characteristics see Extended Data Table 1.

## Blood samples processing and storage

Samples collected at NIH were drawn from participants at the study visit and processed within 24 hours. Briefly, whole blood samples were subjected to Ficoll gradient centrifugation after 1:1 dilution in PBS. Plasma and PBMC samples were obtained through phase separation of plasma layer and Buffy coat phase, respectively. PBMCs were further prepared through centrifugation, red blood cells lysis and washing steps, and stored in CellBanker cell freezing media (Amsbio). All samples were aliquoted and stored at −80 °C and shipped on dry ice. Prior to experiments, aliquots of plasma samples were heat-inactivated (56°C for 1 hour) and then stored at 4°C. Peripheral Blood Mononuclear Cells (PBMCs) obtained from samples collected at Rockefeller University were purified as previously reported^7,8^ by gradient centrifugation and stored in liquid nitrogen in the presence of FCS and DMSO. Heparinized plasma samples were aliquoted and stored at −20°C or less. Prior to experiments, aliquots of plasma samples were heat-inactivated (56°C for 1 hour) and then stored at 4°C.

## ELISAs

ELISAs^51,52^ to evaluate antibodies binding to SARS-CoV-2 S (BioHub), RBD and additional mutated RBDs were performed by coating of high-binding 96-half-well plates (Corning 3690) with 50 μl per well of a 1μg/ml protein solution in PBS overnight at 4 °C. Plates were washed 6 times with washing buffer (1× PBS with 0.05% Tween-20 (Sigma-Aldrich)) and incubated with 170 μl per well blocking buffer (1× PBS with 2% BSA and 0.05% Tween-20 (Sigma)) for 1 h at room temperature. Immediately after blocking, monoclonal antibodies or plasma samples were added in PBS and incubated for 1 h at room temperature. Plasma samples were assayed at a 1:66.6 (RU samples) or a 1:33.3 (NIH samples) starting dilution and 7 additional threefold serial dilutions. Monoclonal antibodies were tested at 10 μg/ml starting concentration and 10 additional fourfold serial dilutions. Plates were washed 6 times with washing buffer and then incubated with anti-human IgG, IgM or IgA secondary antibody conjugated to horseradish peroxidase (HRP) (Jackson Immuno Research 109-036-088 109-035-129 and Sigma A0295) in blocking buffer at a 1:5,000 dilution (IgM and IgG) or 1:3,000 dilution (IgA). Plates were developed by addition of the HRP substrate, TMB (ThermoFisher) for 10 min (plasma samples) or 4 minutes (monoclonal antibodies), then the developing reaction was stopped by adding 50 μl 1 M H_2_SO_4_ and absorbance was measured at 450 nm with an ELISA microplate reader (FluoStar Omega, BMG Labtech) with Omega and Omega MARS software for analysis. For plasma samples, a positive control (plasma from participant COV72^7,8^, diluted 66.6-fold and with seven additional threefold serial dilutions in PBS) was added to every assay plate for validation. The average of its signal was used for normalization of all of the other values on the same plate with Excel software before calculating the area under the curve using Prism V8.4 (GraphPad). For monoclonal antibodies, the EC50 was determined using four-parameter nonlinear regression (GraphPad Prism V8.4).

## Expression of RBD proteins

Mammalian expression vectors encoding the RBDs of SARS-CoV-2 (GenBank MN985325.1; S protein residues 319-539) and eight additional mutant RBD proteins (E484K, Q493R, R346S, N493K, N440K, V367F, A475V, S477N and V483A) with an N-terminal human IL-2 or Mu phosphatase signal peptide were previously described^30^.

## Cells and viruses

293T_Ace2_ ^8^, 293T/ACE2.cl22 and HT1080_/_ACE2.cl14 cells^10^ were cultured in Dulbecco’s Modified Eagle Medium (DMEM) supplemented with 10% fetal bovine serum (FBS) at 37°C and 5% CO_2_. Cells were periodically tested for contamination with mycoplasma or retroviruses. rVSV/SARS-CoV-2/GFP chimeric virus stocks were generated by infecting 293T/ACE2.cl22 cells. Supernatant was harvested 1 day post infection (dpi), cleared of cellular debris, and stored at −80°C. A plaque purified variant designated rVSV/SARS-CoV-2/GFP_2E1_ that encodes D215G/R683G substitutions was used in these studies^10^.

## Selection and analysis of antibody escape mutations

For the selection of monoclonal antibody-resistant spike variants, an rVSV/SARS-CoV-2/GFP_2E1_ (for details see^10^) population containing 10^6^ infectious units was incubated with monoclonal antibodies at 10-40 μg/ml for 1 hr at 37 °C. The virus-antibody mixtures were subsequently incubated with 5 × 10^5^ 293T/ACE2cl.22 cells in 6-well plates. One day after infection the media was replaced with fresh media containing the equivalent concentration of antibodies. Supernatant was harvested 2 days after infection and 150 μl of the cleared supernatant was used to infect cells for passage 2, while 150 μl was subjected to RNA extraction and sequencing.

For identification of putative antibody resistance mutations, RNA was extracted using NucleoSpin 96 Virus Core Kit (Macherey-Nagel). The RNA was reversed transcribed using the SuperScript VILO cDNA Synthesis Kit (Thermo Fisher Scientific). KOD Xtreme Hot Start DNA Polymerase (Millipore Sigma) was used for amplification of cDNA using primers flanking the S-encoding sequence. The PCR products were purified and sequenced as previously described^7,12^. Briefly, tagmentation reactions were performed using 1ul diluted cDNA, 0.25 *μ*l Nextera TDE1 Tagment DNA enzyme (catalog no. 15027865), and 1.25 *μ*l TD Tagment DNA buffer (catalog no. 15027866; Illumina). Next, the DNA was ligated to unique i5/i7 barcoded primer combinations using the Illumina Nextera XT Index Kit v2 and KAPA HiFi HotStart ReadyMix (2X; KAPA Biosystems) and purified using AmPure Beads XP (Agencourt), after which the samples were pooled into one library and subjected to paired-end sequencing using Illumina MiSeq Nano 300 V2 cycle kits (Illumina) at a concentration of 12pM.

For analysis of the sequencing data, the raw paired-end reads were pre-processed to remove trim adapter sequences and to remove low-quality reads (Phred quality score < 20) using BBDuk. Reads were mapped to the rVSV/SARS-CoV-2/GFP sequence using Geneious Prime (Version 2020.1.2). Mutations were annotated using Geneious Prime, with a P-value cutoff of 10^−6^. Because reads have randomly generated ends and different lengths, the mutations do not necessarily have to occur on the same read. e.g K417N and N501Y might occur on the same read or on different reads. The percentages calculated for position X are calculated based on all the reads, and not just the reads, that include position X.

## SARS-CoV-2 pseudotyped reporter virus

A panel of plasmids expressing RBD-mutant SARS-CoV-2 spike proteins in the context of pSARS-CoV-2-S _Δ19_ (based on (NC_045512)) have been described previously^12^. Additional substitutions were introduced using either PCR primer-mediated mutagenesis or with synthetic gene fragments (IDT) followed by Gibson assembly. The mutants E484K and KEN (K417N+E484K+N501Y) were constructed in the context of a pSARS-CoV-2-S _Δ19_ variant with a mutation in the furin cleavage site (R683G). The NT50s and IC50 of these pseudotypes were compared to a wildtype SARS-CoV-2 spike sequence carrying R683G in the subsequent analyses, as appropriate.

Generation of SARS-CoV-2 pseudotyped HIV-1 particles was performed as previously described^8^. Briefly, 293T cells were transfected with pNL4-3ΔEnv-nanoluc and pSARS-CoV-2-S_Δ19_ and pseudotyped virus stocks were harvested 48 hours after transfection, filtered and stored at −80℃.

## SARS-CoV-2 pseudotype neutralization assays

Plasma or monoclonal antibodies from vaccine recipients were four-fold or five-fold serially diluted and then incubated with SARS-CoV-2 pseudotyped HIV-1 reporter virus for 1 h at 37 °C. The antibody and pseudotyped virus mixture was added to 293T_Ace2_ cells^8^ (for comparisons of plasma or monoclonal antibodies from COVID-19-convalescents and vaccine recipients) or HT1080ACE2.cl14 cells^10^ (for analysis of spike mutants with vaccine recipient plasma or monoclonal antibodies). After 48 h cells were washed with PBS and lysed with Luciferase Cell Culture Lysis 5× reagent (Promega) and Nanoluc Luciferase activity in lysates was measured using the Nano-Glo Luciferase Assay System (Promega) with the Glomax Navigator (Promega). The relative luminescence units were normalized to those derived from cells infected with SARS-CoV-2 pseudotyped virus in the absence of plasma or monoclonal antibodies. The half-maximal and 80% or 90% neutralization titers for plasma (NT_50_ and NT_80_/NT_90_, respectively) or half-maximal and 90% inhibitory concentrations for monoclonal antibodies (IC_50_ and IC_90_, respectively) were determined using four-parameter nonlinear regression (least squares regression method without weighting; constraints: top=1, bottom=0) (GraphPad Prism).

## Biotinylation of viral protein for use in flow cytometry

Purified and Avi-tagged SARS-CoV-2 RBD was biotinylated using the Biotin-Protein Ligase-BIRA kit according to manufacturer’s instructions (Avidity) as described before^8^. Ovalbumin (Sigma, A5503-1G) was biotinylated using the EZ-Link Sulfo-NHS-LC-Biotinylation kit according to the manufacturer’s instructions (Thermo Scientific). Biotinylated ovalbumin was conjugated to streptavidin-BV711 (BD biosciences, 563262) and RBD to streptavidin-PE (BD Biosciences, 554061) and streptavidin-AF647 (Biolegend, 405237)^8^.

## Flow cytometry and single cell sorting

Single-cell sorting by flow cytometry was performed as described previously^8^. Briefly, peripheral blood mononuclear cells were enriched for B cells by negative selection using a pan-B-cell isolation kit according to the manufacturer’s instructions (Miltenyi Biotec, 130-101-638). The enriched B cells were incubated in FACS buffer (1× PBS, 2% FCS, 1 mM EDTA) with the following anti-human antibodies (all at 1:200 dilution): anti-CD20-PECy7 (BD Biosciences, 335793), anti-CD3-APC-eFluro 780 (Invitrogen, 47-0037-41), anti-CD8-APC-eFluor 780 (Invitrogen, 47-0086-42), anti-CD16-APC-eFluor 780 (Invitrogen, 47-0168-41), anti-CD14-APC-eFluor 780 (Invitrogen, 47-0149-42), as well as Zombie NIR (BioLegend, 423105) and fluorophore-labelled RBD and ovalbumin (Ova) for 30 min on ice. Single CD3−CD8−CD14−CD16−CD20+Ova−RBD-PE+RBD-AF647+ B cells were sorted into individual wells of 96-well plates containing 4 μl of lysis buffer (0.5× PBS, 10 mM DTT, 3,000 units/ml RNasin Ribonuclease Inhibitors (Promega, N2615) per well using a FACS Aria III and FACSDiva software (Becton Dickinson) for acquisition and FlowJo for analysis. The sorted cells were frozen on dry ice, and then stored at −80°C or immediately used for subsequent RNA reverse transcription.

## Antibody sequencing, cloning and expression

Antibodies were identified and sequenced as described previously^8^. In brief, RNA from single cells was reverse-transcribed (SuperScript III Reverse Transcriptase, Invitrogen, 18080-044) and the cDNA stored at −20°C or used for subsequent amplification of the variable IGH, IGL and IGK genes by nested PCR and Sanger sequencing. Sequence analysis was performed using MacVector. Amplicons from the first PCR reaction were used as templates for sequence- and ligation-independent cloning into antibody expression vectors. Recombinant monoclonal antibodies and Fabs were produced and purified as previously described^8^.

## Cryo-EM sample preparation

Expression and purification of SARS-CoV-2 6P stabilized S trimers^37^ was conducted as previously described^53^. Purified Fab and S 6P trimer were incubated at a 1.1:1 molar ratio per protomer on ice for 30 minutes prior to deposition on a freshly glow-discharged 300 mesh, 1.2/1.3 Quantifoil copper grid. Immediately before 3 μl of complex was applied to the grid, fluorinated octyl-malotiside was added to the Fab-S complex to a final detergent concentration of 0.02% w/v, resulting in a final complex concentration of 3 mg/ml. Samples were vitrified in 100% liquid ethane using a Mark IV Vitrobot after blotting for 3 s with Whatman No. 1 filter paper at 22°C and 100% humidity.

## Cryo-EM data collection and processing

Data collection and processing followed a similar workflow to what has been previously described in detail^29^. Briefly, micrographs were collected on a Talos Arctica transmission electron microscope (Thermo Fisher) operating at 200 kV for all Fab-S complexes. Data were collected using SerialEM automated data collection software^54^ and movies were recorded with a K3 camera (Gatan). For all datasets, cryo-EM movies were patch motion corrected for beam-induced motion including dose-weighting within cryoSPARC v2.15^55^ after binning super resolution movies. The non-dose-weighted images were used to estimate CTF parameters using cryoSPARC implementation of the Patch CTF job. Particles were picked using Blob picker and extracted 4× binned and 2D classified. Class averages corresponding to distinct views with secondary structure features were chosen and ab initio models were generated. 3D classes that showed features of a Fab-S complex were re-extracted, unbinned (0.869 Å/pixel) and homogenously refined with C1 symmetry. Overall resolutions were reported based on gold standard FSC calculations.

## Cryo-EM Structure Modeling and Refinement

Coordinates for initial complexes were generated by docking individual chains from reference structures into cryo-EM density using UCSF Chimera^56^ (S trimer: PDB 6KXL, Fab: PDB 6XCA or 7K8P after trimming CDR3 loops and converting to a polyalanine model). Models were then refined into cryo-EM maps by rigid body and real space refinement in Phenix^57^. If the resolution allowed, partial CDR3 loops were built manually in Coot^58^ and then refined using real-space refinement in Phenix.

## Computational analyses of antibody sequences

Antibody sequences were trimmed based on quality and annotated using Igblastn v.1.14. with IMGT domain delineation system. Annotation was performed systematically using Change-O toolkit v.0.4.540 ^59^. Heavy and light chains derived from the same cell were paired, and clonotypes were assigned based on their V and J genes using in-house R and Perl scripts (Fig. 2c and f, Extended Data Fig. 3d, Extended Data Fig. 4b). All scripts and the data used to process antibody sequences are publicly available on GitHub (https://github.com/stratust/igpipeline).

The frequency distributions of human V genes in anti-SARS-CoV-2 antibodies from this study was compared to 131,284,220 IgH and IgL sequences generated by ^60^ and downloaded from cAb-Rep ^61^, a database of human shared BCR clonotypes available at https://cab-rep.c2b2.columbia.edu/. Based on the 97 distinct V genes that make up the 4186 analyzed sequences from Ig repertoire of the 14 participants present in this study, we selected the IgH and IgL sequences from the database that are partially coded by the same V genes and counted them according to the constant region. The frequencies shown in (Fig 2e and Extended Data Fig 4a) are relative to the source and isotype analyzed. We used the two-sided binomial test to check whether the number of sequences belonging to a specific IgHV or IgLV gene in the repertoire is different according to the frequency of the same IgV gene in the database. Adjusted p-values were calculated using the false discovery rate (FDR) correction. Significant differences are denoted with stars.

Nucleotide somatic hypermutation and CDR3 length were determined using in-house R and Perl scripts. For somatic hypermutations, IGHV and IGLV nucleotide sequences were aligned against their closest germlines using Igblastn and the number of differences were considered nucleotide mutations. The average mutations for V genes were calculated by dividing the sum of all nucleotide mutations across all participants by the number of sequences used for the analysis. To calculate the GRAVY scores of hydrophobicity^62^ we used Guy H.R. Hydrophobicity scale based on free energy of transfer (kcal/mole)^63^ implemented by the R package Peptides (the Comprehensive R Archive Network repository; https://journal.r-project.org/archive/2015/RJ-2015-001/RJ-2015-001.pdf). We used 1405 heavy chain CDR3 amino acid sequences from this study and 22,654,256 IGH CDR3 sequences from the public database of memory B cell receptor sequences^64^. The two-tailed Wilcoxon nonparametric test was used to test whether there is a difference in hydrophobicity distribution.

## Data availability statement

Data are provided in Extended Data Tables 1-7. The raw sequencing data and computer scripts associated with Fig. 2 have been deposited at Github (https://github.com/stratust/igpipeline). This study also uses data from “A Public Database of Memory and Naive B-Cell Receptor Sequences” ^64^, PDB (6VYB and 6NB6) and from “High frequency of shared clonotypes in human B cell receptor repertoires” ^60^. Cryo-EM maps associated with data reported in this manuscript will be deposited in the Electron Microscopy Data Bank (EMDB: https://www.ebi.ac.uk/pdbe/emdb/) under accession codes EMD-23393 (C601-S), EMD-23394 (C603-S), EMD-23395 (C643-S), EMD-23396 (C663-S), EMD-23397 (C666-S), EMD-23398 (C669-S), and EMD-23399 (C670-S).

## Data presentation

Figures were arranged in Adobe Illustrator 2020.

## Code availability statement

Computer code to process the antibody sequences is available at GitHub (https://github.com/stratust/igpipeline).

## Acknowledgements

We thank all study participants who devoted time to our research; Drs. Barry Coller and Sarah Schlesinger, the Rockefeller University Hospital Clinical Research Support Office and nursing staff; Charles M. Rice and all members of the M.C.N. laboratory for helpful discussions and Maša Jankovic for laboratory support; and Dr. Jost Vielmetter and the Protein Expression Center in the Beckman Institute at Caltech for expression assistance. Electron microscopy was performed in the Caltech Beckman Institute Resource Center for Transmission Electron Microscopy and we thank Drs. Songye Chen and Andrey Malyutin for technical assistance. This work was supported by NIH grant P01-AI138398-S1 (M.C.N. and P.J.B.) and 2U19AI111825 (M.C.N.); the Caltech Merkin Institute for Translational Research and P50 AI150464-13 (P.J.B.), a George Mason University Fast Grant (P.J.B.); R37-AI64003 to P.D.B.; R01AI78788 to T.H.; We thank Dr. Jost Vielmetter and the Protein Expression Center in the Beckman Institute at Caltech for expression assistance C.O.B. is supported by the HHMI Hanna Gray and Burroughs Wellcome PDEP fellowships. C.G. was supported by the Robert S. Wennett Post-Doctoral Fellowship, in part by the National Center for Advancing Translational Sciences (National Institutes of Health Clinical and Translational Science Award program, grant UL1 TR001866), and by the Shapiro-Silverberg Fund for the Advancement of Translational Research. P.D.B. and M.C.N. are Howard Hughes Medical Institute Investigators.

## Author Contributions

P.D.B., P.J.B., R.C., T.H., M.C.N, Z.W., F.S., Y.W. F.M. C.O.B, S.F., D.S.B., M.Cipolla. conceived, designed and analyzed the experiments. M.Caskey, C.G., J.A. L, K.W. D.S.B. designed clinical protocols Z.W., F.S., Y.W. F.M. C.O.B, S.F., D.S.B., M.Cipolla, J.D.S. A.G. Z.Y., M.E.A., K.E.H. carried out experiments. C.G., M.Caskey, K.W. D., R.A.C., A.H., K.G.M. recruited participants and executed clinical protocols. I.S., R.P, J.D., J.X. and C.U.O. processed clinical samples. T.Y.O. and V.R. performed bioinformatic analysis. R.C. P.D.B., P.J.B., T.H., and M.C.N. wrote the manuscript with input from all co-authors.

---

**Extended Data Fig. 1.**
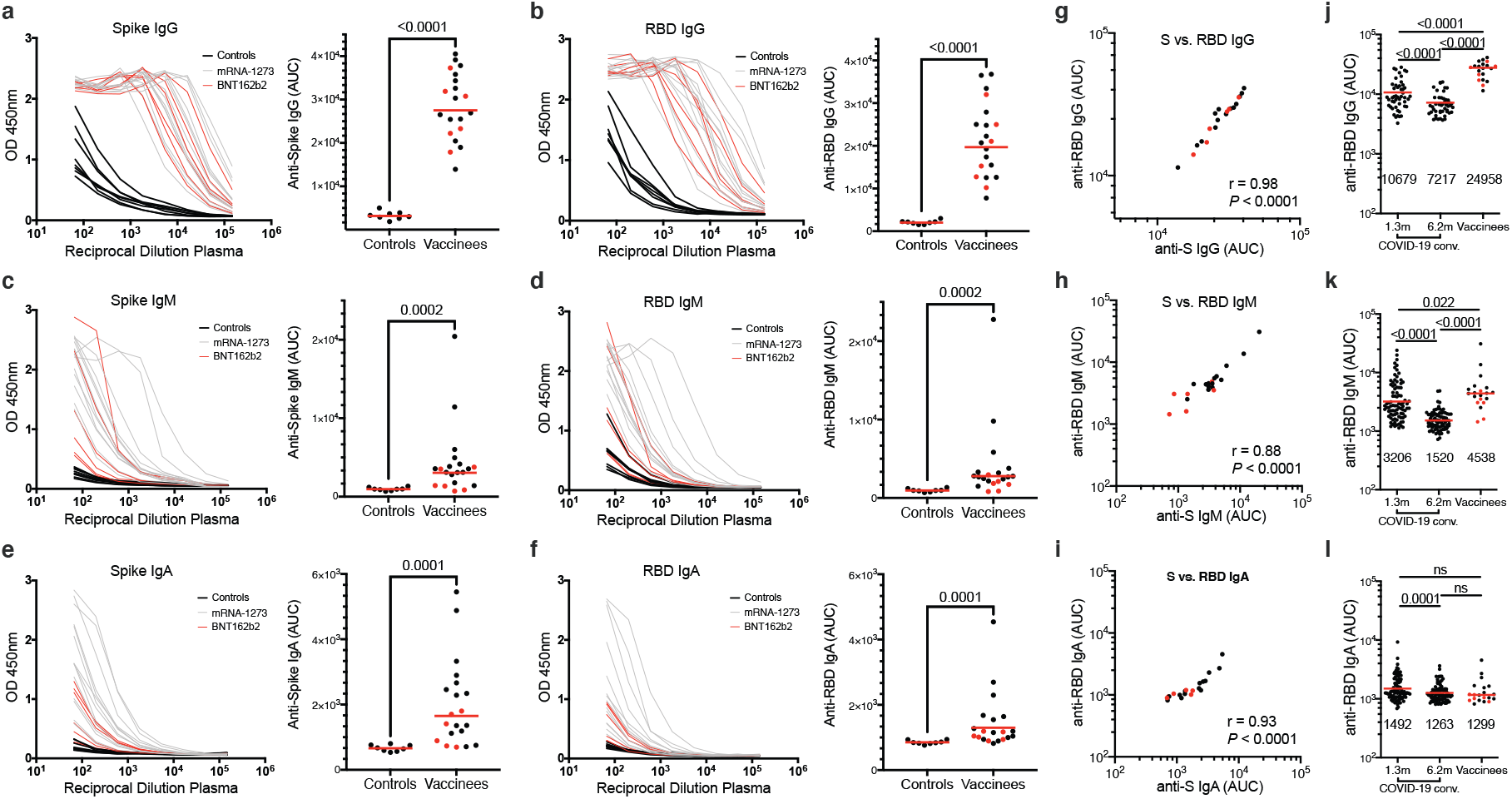
Plasma antibodies against SARS-CoV-2. **a**–**f**, Results of ELISAs measuring plasma reactivity to S (**a**,**c,e**) and RBD protein (**b,d,f**) of 20 vaccinees (grey curves) and 8 controls (black curves). **a**, Anti-S IgG. **b**, Anti-RBD IgG. **c**, Anti-S IgM. **d**, Anti-RBD IgM. **e**, Anti-S IgA. **f**, Anti-RBD IgA. Left, optical density at 450 nm (OD 450 nm) for the indicated reciprocal plasma dilutions. Right, normalized area under the curve (AUC) values for the 8 controls and 20 vaccinees. Horizontal bars indicate geometric mean. Statistical significance was determined using the two-tailed Mann–Whitney U-test. Average of two or more experiments. **g-i,** Correlations of plasma antibodies measurements. **g,** Normalized AUC for IgG anti-S (X axis) plotted against normalized AUC for IgG anti-RBD (Y axis). **h,** Normalized AUC for IgM anti-S (X axis) plotted against normalized AUC for IgM anti-RBD (Y axis). **i,** Normalized AUC for IgA anti-S (X axis) plotted against normalized AUC for IgA anti-RBD (Y axis). The *r* and *p* values in **g-i** were determined with the two-tailed Spearman’s correlation test. Moderna vaccinees are in black and Pfizer-BioNTech in red. **j-l,** Results of ELISAs measuring plasma reactivity to RBD in convalescent volunteers 1.3 and 6.2 months after infection^7,8^ and in 20 vaccinees, who received the Moderna vaccine (black dots) and Pfizer-BioNTech vaccine (red dots). **j,** Anti-RBD IgG. **k,** Anti-RBD IgM. **l,** Anti-RBD IgA. The normalized area under the curve (AUC) values are shown. Positive and negative controls were included for validation. Red horizontal bars and indicated values represent geometric mean. Statistical significance was determined using two-tailed Mann–Whitney U-test or Wilcoxon matched-pairs signed rank test.

**Extended Data Fig. 2:**
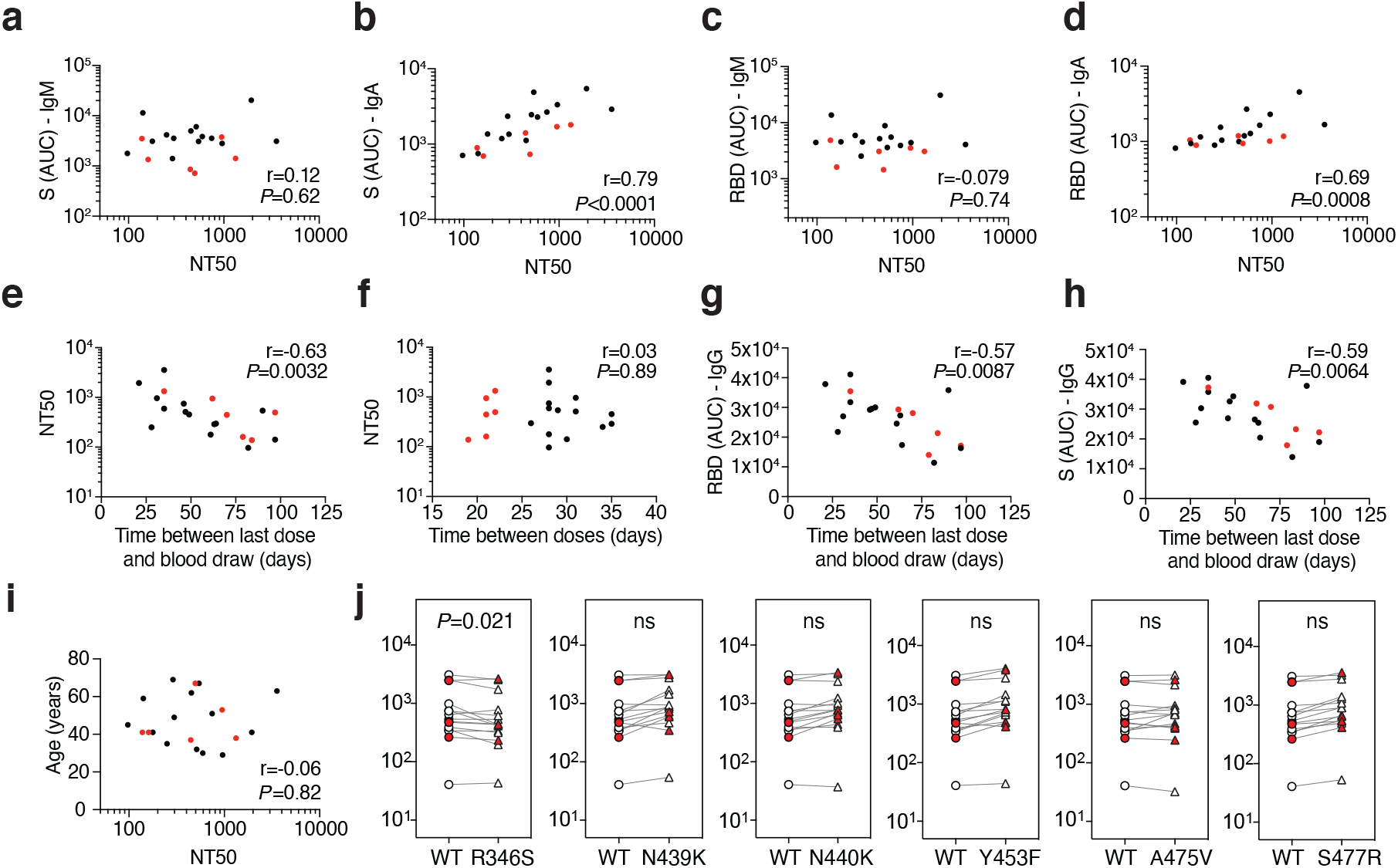
Plasma neutralizing activity. **a**, Anti-S IgM AUC (Y axis) plotted against NT_50_ (X axis) r=0.12, p<0.62. **b,** Anti-S IgA AUC (Y axis) plotted against NT_50_ (X axis) r=0.79, p<0.0001. **c**, Anti-RBD IgM AUC (Y axis) plotted against NT50 (X axis) r=−0.079 p=0.74. **d**, Anti-RBD IgA AUC (Y axis) plotted against NT50 (X axis) r=0.69 p=0.0008. **e**, NT_50_ (Y axis) plotted against time between last dose and blood draw (X axis) r=−0.63 p=0.0032. **f**, NT_50_ (Y axis) plotted against time between doses (X axis) r=0.03 p=0.89. **g**, Anti-RBD IgG AUC (Y axis) plotted against time between last dose and blood draw (X axis) r=−0.57 p=0.0084. **h**, Anti-S IgG AUC (Y axis) plotted against time between last dose and blood draw (X axis) r=−0.59 p=0.0064. **i**, Age (Y axis) plotted against NT_50_ (X axis) r=−0.06 p=0.82. The r and p values were determined by two-tailed Spearman’s. Moderna vaccinees in black and Pfizer-BioNTech in red. **j**, NT50 values for vaccinee plasma (n=15) neutralization of pseudotyped viruses with WT and the indicated RBD-mutant SARS-CoV-2 S proteins; p-values determined using one tailed t-test.

**Extended Data Fig. 3:**
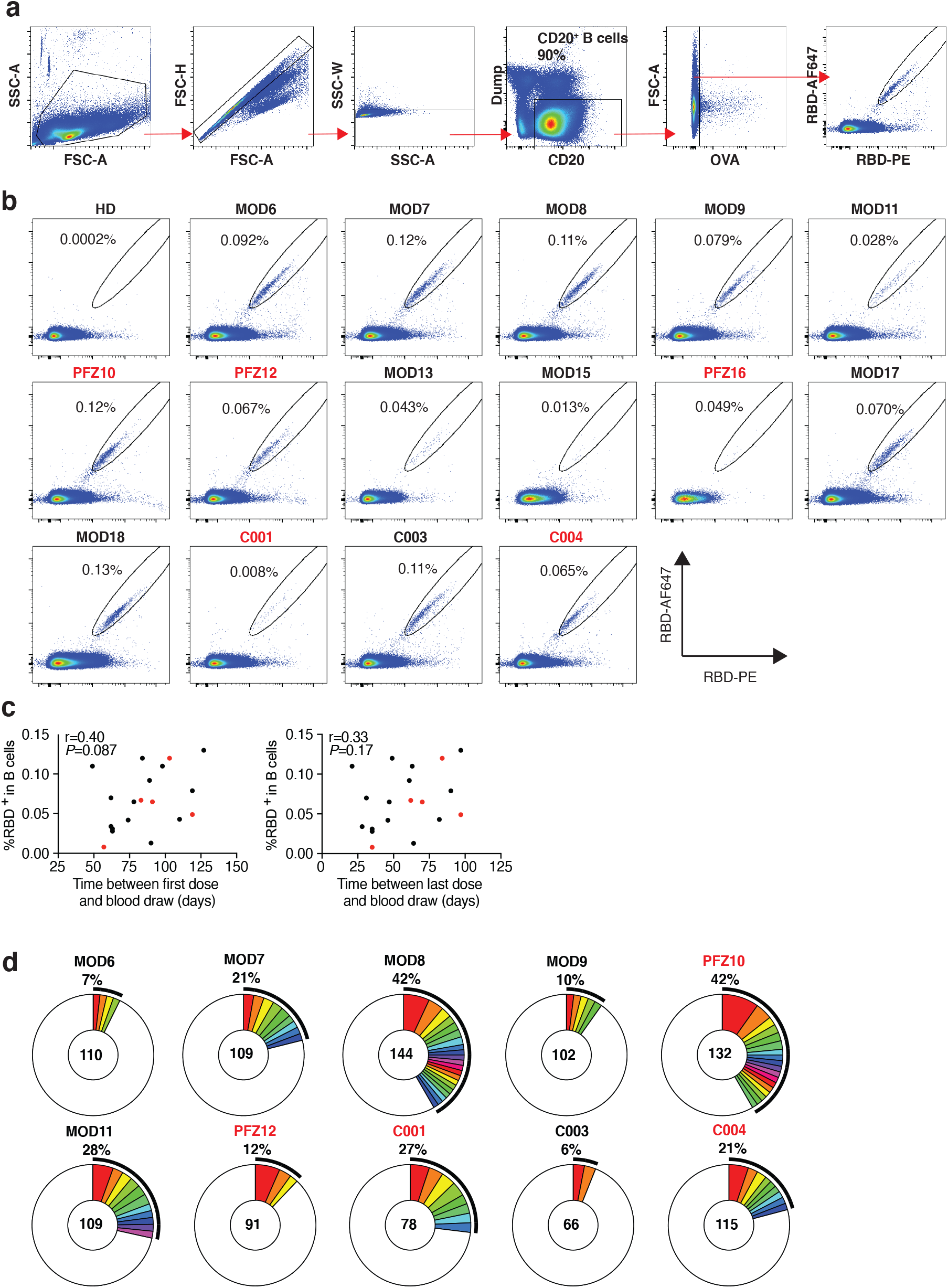
Flow cytometry. **a,** Gating strategy used for cell sorting. Gating was on singlets that were CD20+ and CD3-CD8-CD14-CD16-OVA-. Sorted cells were RBD-PE+ and RBD-AF647+. **b,** Flow cytometry showing the percentage of RBD-double positive memory B cells from a pre-COVID-19 control (HD) and 15 vaccinees, who received the Moderna vaccine are shown in black and Pfizer-BioNTech vaccine recipients are in red. **c**, the percentage of RBD-binding memory B cells in vaccinees (Y axis) plotted against time between first dose and blood draw (X axis) r=0.40 p=0.087 (left panel), and between last dose and blood draw (X axis) r=0.33 p=0.17 (right panel). Moderna vaccinees in black and Pfizer-BioNTech in red. The r and p values for correlations were determined by two-tailed Spearman’s. **d,** Pie charts show the distribution of antibody sequences from 10 individuals in **b.** The number in the inner circle indicates the number of sequences analyzed. Pie slice size is proportional to the number of clonally related sequences. The black outline indicates the frequency of clonally expanded sequences. The r and p values for correlations in **c** were determined by the two-tailed Spearman correlation test.

**Extended Data Fig. 4:**
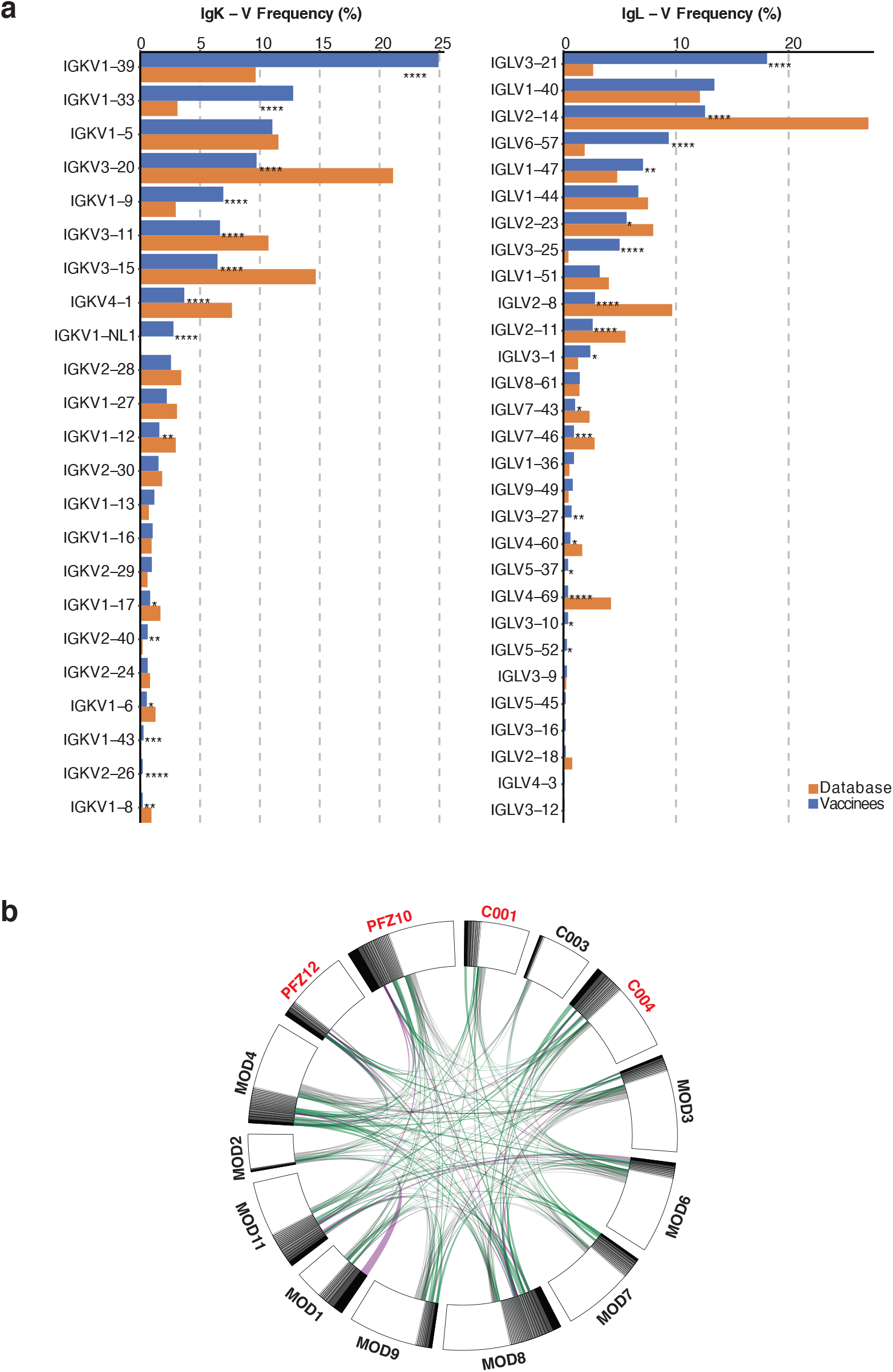
Frequency distributions of human VL genes. Graph shows relative abundance of human IGVK (left) and IgVL (right) genes of Sequence Read Archive accession SRP010970 (orange)^65^, and vaccinees (blue). Two-sided binomial tests with unequal variance were used to compare the frequency distributions., significant differences are denoted with stars (* p < 0.05, ** p < 0.01, *** p < 0.001, **** = p < 0.0001). **b.** Sequences from 14 individuals (Extended Data Table 3) with clonal relationships. Interconnecting lines indicate the relationship between antibodies that share V and J gene segment sequences at both IGH and IGL. Purple, green and grey lines connect related clones, clones and singles, and singles to each other, respectively.

**Extended Data Fig. 5:**
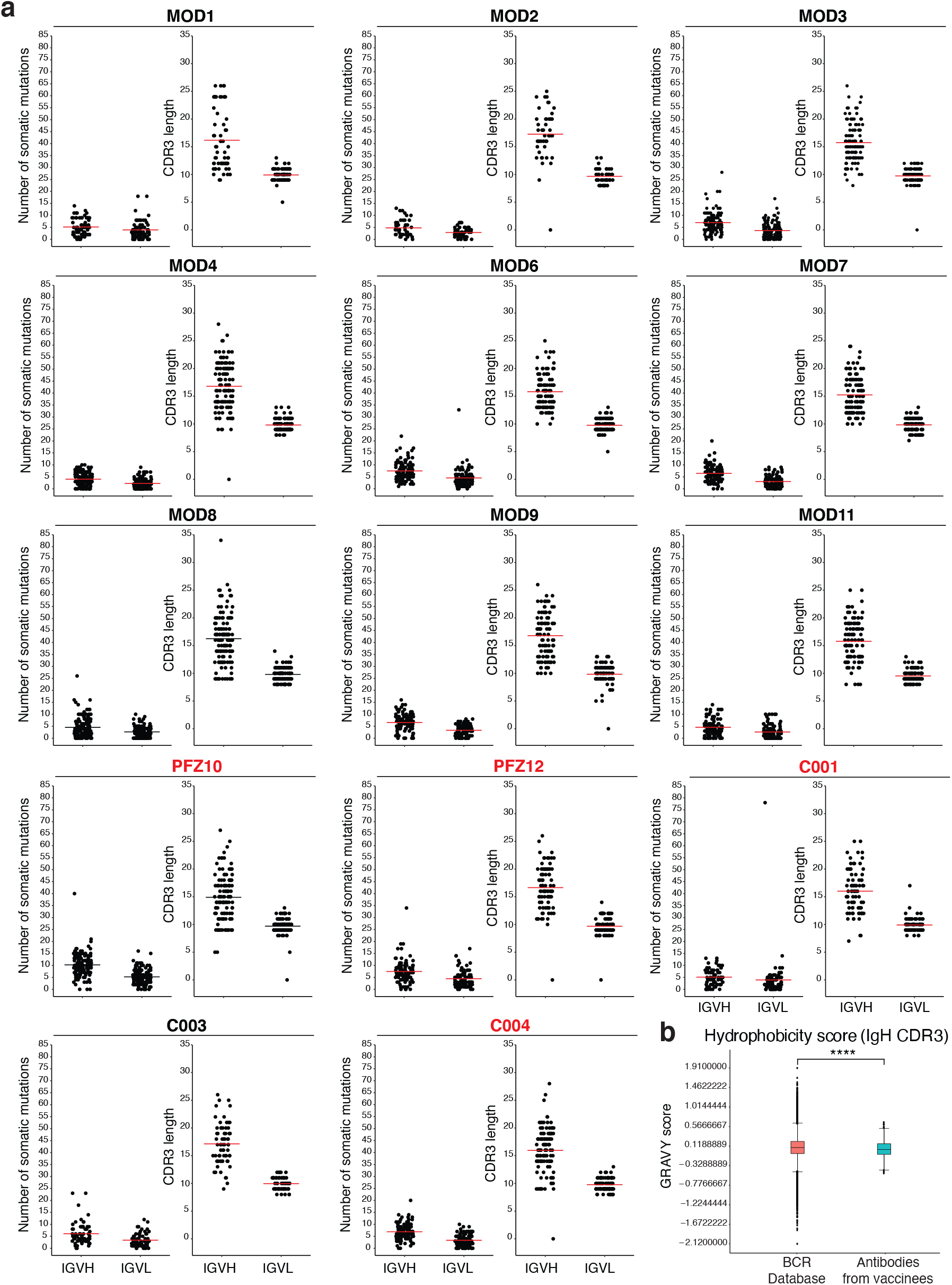
Antibody somatic hypermutation, and CDR3 length. **a,** Number of somatic nucleotide mutations in both the IGVH and IGVL in 14 participants (left). Individuals who received the Moderna vaccine are shown in black and Pfizer-BioNTech vaccine recipients in red. For each individual, the number of the amino acid length of the CDR3s at the IGVH and IGVL is shown (right). The horizontal bars indicate the mean. The number of antibody sequences (IGVH and IGVL) evaluated for each participant are n=68 (MOD1), n=45 (MOD2), n=117 (MOD3), n=123 (MOD4), n=110 (MOD6), n=109 (MOD7), n=144 (MOD8), n=102 (MOD9), n=132 (PFZ10), n=109 (MOD11), n=91 (PFZ12), n=78 (C001), n=66 (C003), and n=115 (C004). **b,** Distribution of the hydrophobicity GRAVY scores at the IGH CDR3 compared to a public database (see Methods for statistical analysis). The box limits are at the lower and upper quartiles, the center line indicates the median, the whiskers are 1.5× interquartile range and the dots represent outliers. Statistical significance was determined using two-tailed Wilcoxon matched-pairs signed rank test (**** = p < 0.0001).

**Extended Data Fig. 6:**
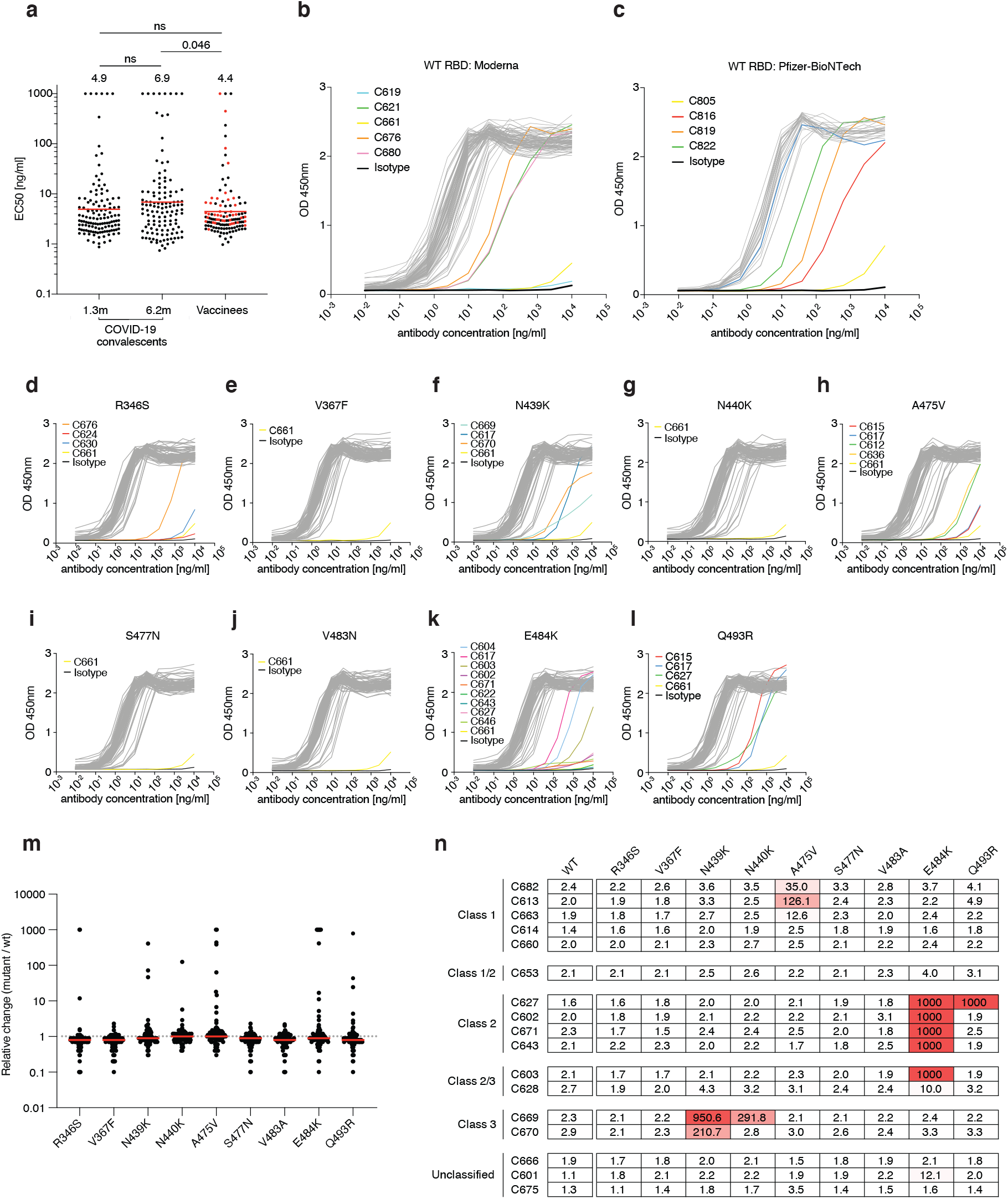
Monoclonal antibody ELISAs. **a,** Graphs show anti-SARS-CoV-2 RBD antibody reactivity. ELISA EC50 values for all antibodies isolated from COVID-19 convalescent individuals assayed at 1.3 and 6.2 months after infection^7,8^ and 127 selected monoclonal antibodies isolated from 4 Moderna vaccinees (black dots) and 4 Pfizer-BioNTech vaccinees (red dots) measured at 8 weeks after the boost. Red horizontal bars and indicated values represent geometric mean. Statistical significance was determined using two-tailed Mann–Whitney U-test. **b-c**, Graphs show ELISA titration curves for 86 monoclonal antibodies isolated from Moderna vaccinees (**b**) and 41 monoclonal antibodies isolated from Pfizer-BioNTech vaccinees (**c**). **d-l,** Graphs show ELISA titrations for 84 antibodies isolated from Moderna vaccinees against the indicated RBD variants. Isotype control and low-binding antibodies are indicated in colors. C661 is a non-binding antibody. Data are representative of two independent experiments. **m,** Relative change in EC50 values for the indicated RBD variants over wt RBD of 84 antibodies isolated from Moderna vaccinees. Red horizontal bars represent geometric mean. **n,** a heat map summary of EC_50_ values for binding to wild type RBD and the indicated mutant RBDs for 17 top neutralizing antibodies.

**Extended Data Fig. 7:**
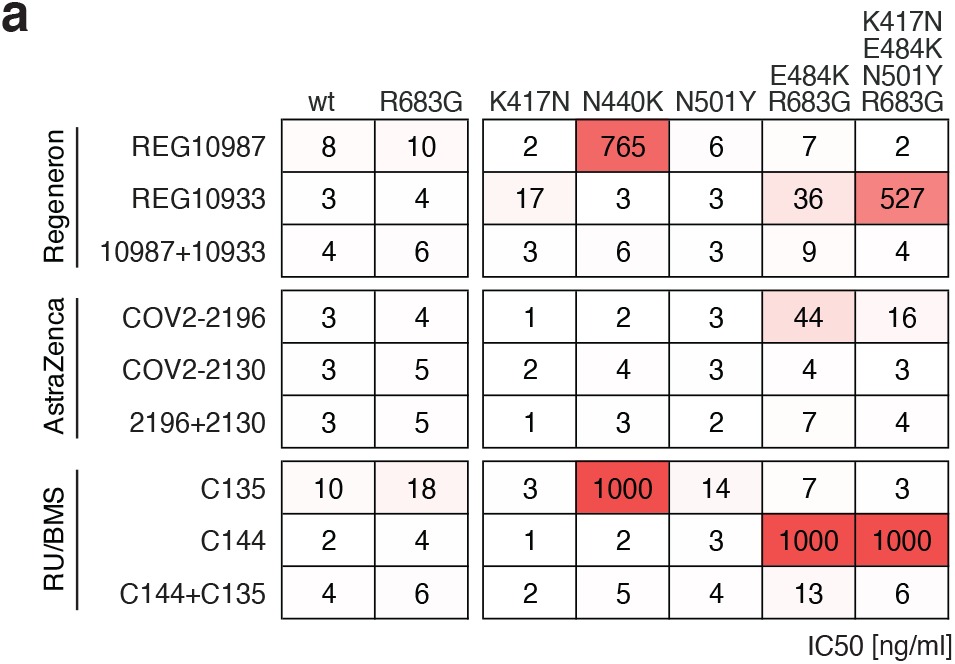
Neutralizing activity of monoclonal antibodies in clinical development against SARS-CoV-2 variants. **a,** Results of a SARS-CoV-2 pseudovirus neutralization assay. IC_50_ values for 6 different monoclonal antibodies, alone or in their clinically designated combinations, for neutralization of wild type and the indicated mutant SARS-CoV-2 pseudotyped viruses. Antibodies with IC_50_ values above 1000 ng/ml were plotted at 1000 ng/ml. Data are the mean of 2 independent experiments. Color gradient indicates IC_50_ values ranging from 0 (white) to 1000 ng/ml (red). The combination of REGN 10987 and 10933 (casirivimab and imdevimab, respectively)^13,66,67^ has been granted emergency use authorization by the U.S. FDA, the combination of COV2-2196 and COV2-2130 (licensed to Astra Zeneca as AZD7442)^26^, and the combination of C135 and C144 (The Rockefeller University)^8^ are currently in clinical trials (NCT04507256 and NCT04700163, respectively).

**Extended Data Fig. 8:**
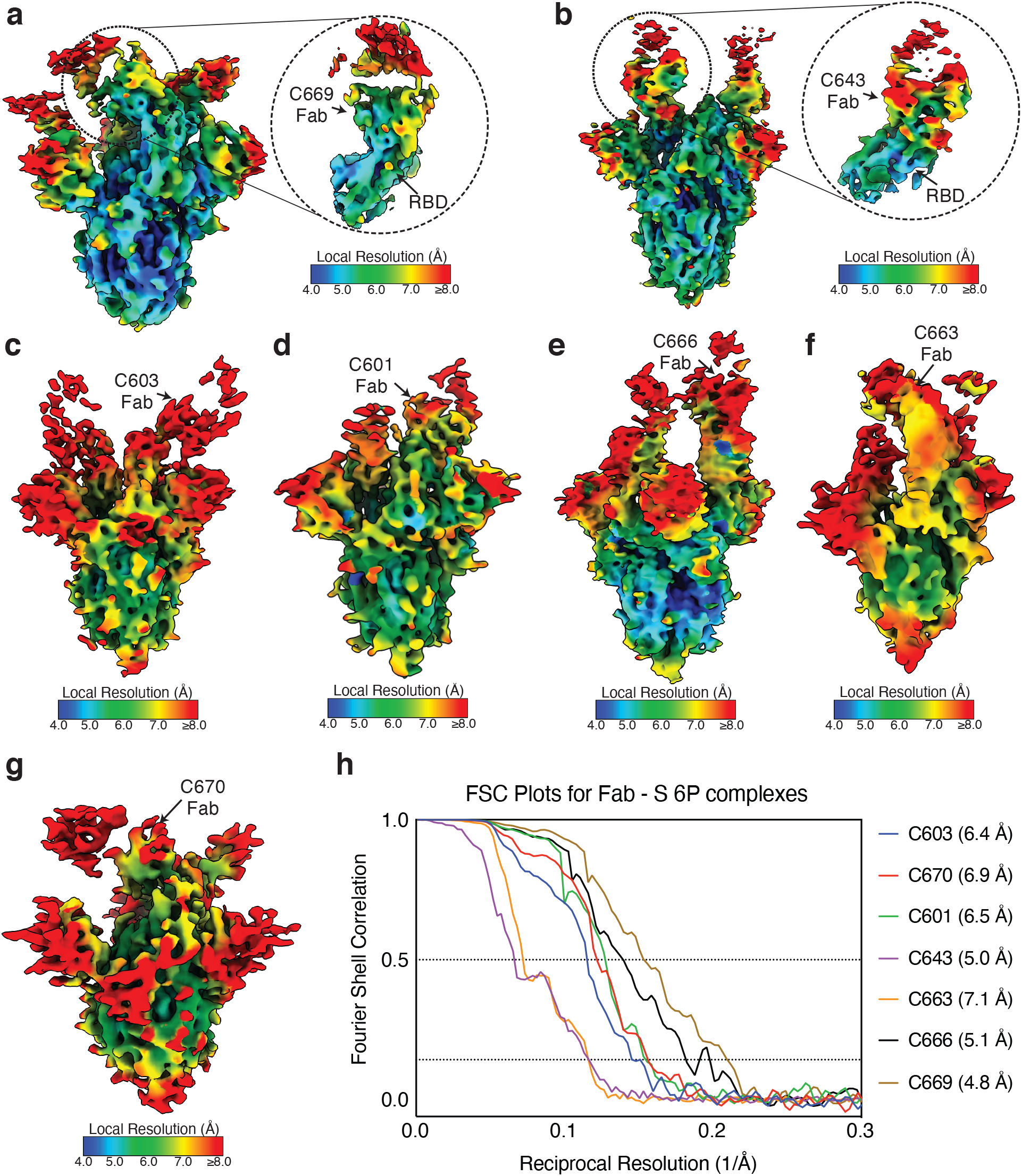
Local resolution estimates of Fab-S cryo-EM reconstructions. **a-g,** Local resolution maps calculated using cryoSPARC for (**a**) C669-S, (**b**) C643-S, (**c**) C603-S, (**d**) C601-S, (**e**) C666-S, (**f**) C663-S, and (**g**) C670-S complexes. Close-up views for Fab-RBD interfaces are highlighted for (**a**) C669 and (**b**) C643. **h,** Gold-standard Fourier shell correlation curves for Fab-S complexes. The 0.5 and 0.143 cutoffs are indicated by dashed lines.

## Supplementary Guide

**<u>Supplementary Table 1:</u> Individual vaccinee characteristics**

**<u>Supplementary Table 2:</u> Neutralization of WT/KEN pseudovirus by convalescent plasma**

**<u>Supplementary Table 3:</u> Antibody sequences from vaccinees is provided as a separate Excel file**

**<u>Supplementary Table 4:</u> CDR3 alignment of highly identical clonal sequences**

**<u>Supplementary Table 5:</u> Sequences, half maximal effective concentrations (EC50s) and inhibitory concentrations (IC50s) of the cloned monoclonal antibodies is provided as a separate Excel file**

**<u>Supplementary Table 6:</u> Neutralization activity of mAbs against mutant SARS-CoV-2 pseudoviruses**

**<u>Supplementary Table 7:</u> Cryo-EM data collection and processing statistics**

